# Two-dimensional NMR from a Single Pulse: Reconstructing Heteronuclear 2D spectra via off-resonance decoupling and Deep Neural Networks

**DOI:** 10.1101/2025.10.30.685498

**Authors:** Nihar P Khandave, Veera Mohana Rao Kakita, Vaibhav Kumar Shukla, Kevin Haubrich, Pramodh Vallurupalli, D Flemming Hansen

## Abstract

Nuclear Magnetic Resonance (NMR) spectroscopy is ubiquitous in many areas of science, including chemistry, material science, biophysics and structural biology. Heteronuclear two-dimensional (2D) NMR experiments form the basis of several NMR based investigations, especially those involving biomolecules like proteins. However, in solution, as the molecule of interest becomes larger, transverse relaxation times of the spins of interest become shorter. This makes it difficult to record 2D ^1^H-^13^C or ^1^H-^15^N correlation maps of large protein molecules using standard Fourier-transform based experiments as they contain transfer delays that are long compared to the short relaxation times. Herein, we explore the possibility of leveraging deep neural networks (DNNs) to obtain 2D correlation maps of proteins while eliminating transfer delays from experiments. In this study, we show that 2D methyl ^1^H-^13^C correlation maps can be obtained from experiments containing only a single ^1^H excitation pulse followed by off-resonance ^13^C continuous-wave decoupling. A DNN was successfully trained to reconstruct the 2D ^1^H-^13^C correlation map from datasets recorded with two ^13^C decoupling fields using protein samples enriched with (ILV) ^13^CHD_2_ methyl groups in a ^2^H background. The efficacy of this strategy is demonstrated by using the DNN to reconstruct methyl ^1^H-^13^C correlation maps of the ∼8 kDa FF domain from human HYPA/FBP11, the ∼18 kDa T4 phage lysozyme, as well as on the ∼360 kDa α_7_α_7_ particle from the *Thermoplasma acidophilum* proteasome. This study illustrates the potential for improving the sensitivity and resolution of NMR spectra using new experiments tailored for use with DNNs.

**Significance statement:** Multidimensional heteronuclear solution state nuclear magnetic resonance (NMR) spectroscopy underpins studies of protein dynamics and interactions. However, the experiments have thus far required magnetization-transfer periods and evolution delays, both of which rapidly erode the signal arising from large biomolecules. We introduce here a single-pulse strategy wherein these delays are replaced by off-resonance continuous-wave decoupling so that the two-dimensional correlation map can be reconstructed from the off-resonance decoupling data using a deep neural network. The approach yields high-quality methyl correlation spectra for proteins spanning ∼8–360 kDa. The co-design of NMR experiments along with deep neural networks establishes a general framework to acquire multidimensional NMR spectra using nontraditional strategies that can be extended to other spin systems and higher-dimensional experiments.

## Introduction

NMR spectroscopy is an important tool amongst others in material science, chemistry and structural biology largely due to the development of Fourier-transform (FT) multidimensional NMR experiments (1). In particular heteronuclear multidimensional NMR experiments have impacted the study of biomolecular structure and dynamics in solution (2-7). Two dimensional (2D) ^1^H-^15^N and ^1^H-^13^C correlation maps are now routinely used to inform on a variety of process, such as conformational dynamics by measuring relaxation properties of nuclei, or biomolecular interactions by monitoring peak positions and linewidths as a function of ligand concentration, etc. (4). The heteronuclear 2D correlation experiments also form the primary building block for most higher dimensional NMR experiments (4, 7).

The key ingredients of the powerful heteronuclear multidimensional experiments are pulse sequence blocks that transfer magnetization between adjacent nuclei via ^1^*J* scalar couplings and chemical shift labeling delays during which the magnetization of interest evolves at the chemical shift of the nucleus being studied (4, 7-9). However, in solution, as the molecules become larger (≳ 100 kDa), transverse relaxation times become shorter, thus leading to both broader peaks as well as reduced signal intensities due to losses during the transfer periods making it difficult to record (even) 2D heteronuclear correlation maps with adequate sensitivity and resolution (4, 10, 11). Hence, different NMR experiments are now being developed to record heteronuclear correlation maps from samples that contain large biomolecules. The strategies that these new experiments employ generally entail (*i*) increasing the transverse relaxation time by evolving coherences with favorable relaxation properties (10-12) or by reducing the ^1^H density via ^2^H enrichment (13) and (*ii*) reducing the length of experiments by utilizing isotope labelling strategies that isolates the spins of interest (14) or reducing the number of the delays in the experiments (15-17).

An upcoming and promising strategy to improve the quality of NMR spectra involves using deep neural networks (DNNs) in combination with new experimental designs, because it is now becoming clear that DNNs can be trained to distil the key information from complex information rich NMR data (18-23). In particular, synergistic approaches that combine the development of NMR pulse-sequences and DNNs are beginning to substantially impact biomolecular NMR. Examples include DNNs that virtually decouple and enhance the resolution of methyl and aromatic ^1^H-^13^C correlation maps acquired on uniformly ^13^C labeled protein samples from relatively large proteins (methyl ∼360 kDa, aromatic ∼50 kDa) that could not be obtained using standard constant time (CT) approaches (24, 25). Aromatic groups in proteins contain ^13^C sites bonded to varying numbers of other ^13^C nuclei leading to singlets, doublets and triplets in the ^13^C dimension of a regular ^1^H-^13^C correlation map. Hence additional information is required to virtually decouple a regular aromatic ^1^H-^13^C spectrum into a resolution enhanced ‘decoupled’ ^1^H-^13^C correlation map that contains only singlets. Providing additional information (along a pseudo-dimension) obtained using a new experiment that encodes the multiplet information allowed the DNN to reconstruct high resolution decoupled ^1^H-^13^C correlation maps, thereby illustrating the power of synergistically developing new NMR experiments along with DNNs (25). Key to the success of this strategy is that additional information is provided (in a pseudo-dimension) to the DNN and that (short) non-CT experiments are recorded, instead of the traditional CT experiments, that are often long compared to the transverse relaxation times of the ^13^C nuclei being studied.

With the ultimate goal of recording heteronuclear 2D correlation maps of large macromolecular systems (preferably uniformly labelled), we sought to decrease the length of the pulse sequences even further than what has been done previously (15, 16). We therefore revisited the idea of using a single excitation pulse followed by off-resonance decoupling to correlate spins that are scalar coupled to one another (26-29). Specifically, herein we explore the possibility of developing a DNN to generate a regular high-resolution 2D ^1^H-^13^C correlation map from these customized data recorded on an isolated ^1^H-^13^C spin-system using an experiment with a single ^1^H excitation pulse (Fig. 1, S1). In this proof of principle study, we find that a simple DNN can transform protein methyl (^13^CHD_2_) off-resonance data into a methyl ^1^H-^13^C correlation map. We demonstrate the applicability of the single pulse method by reconstructing the methyl ^1^H-^13^C correlation maps of three different proteins, that is, the FF domain (30) from human HYPA/FBP11 (FF domain, ∼8 kDa), T4 phage lysozyme (T4L, ∼18 kDa) (31) and the double-*α*-ring particle of the *Thermoplasma acidophilum* proteasome (*α*_7_*α*_7_, ∼360 kDa) (32).

**Fig. 1.**
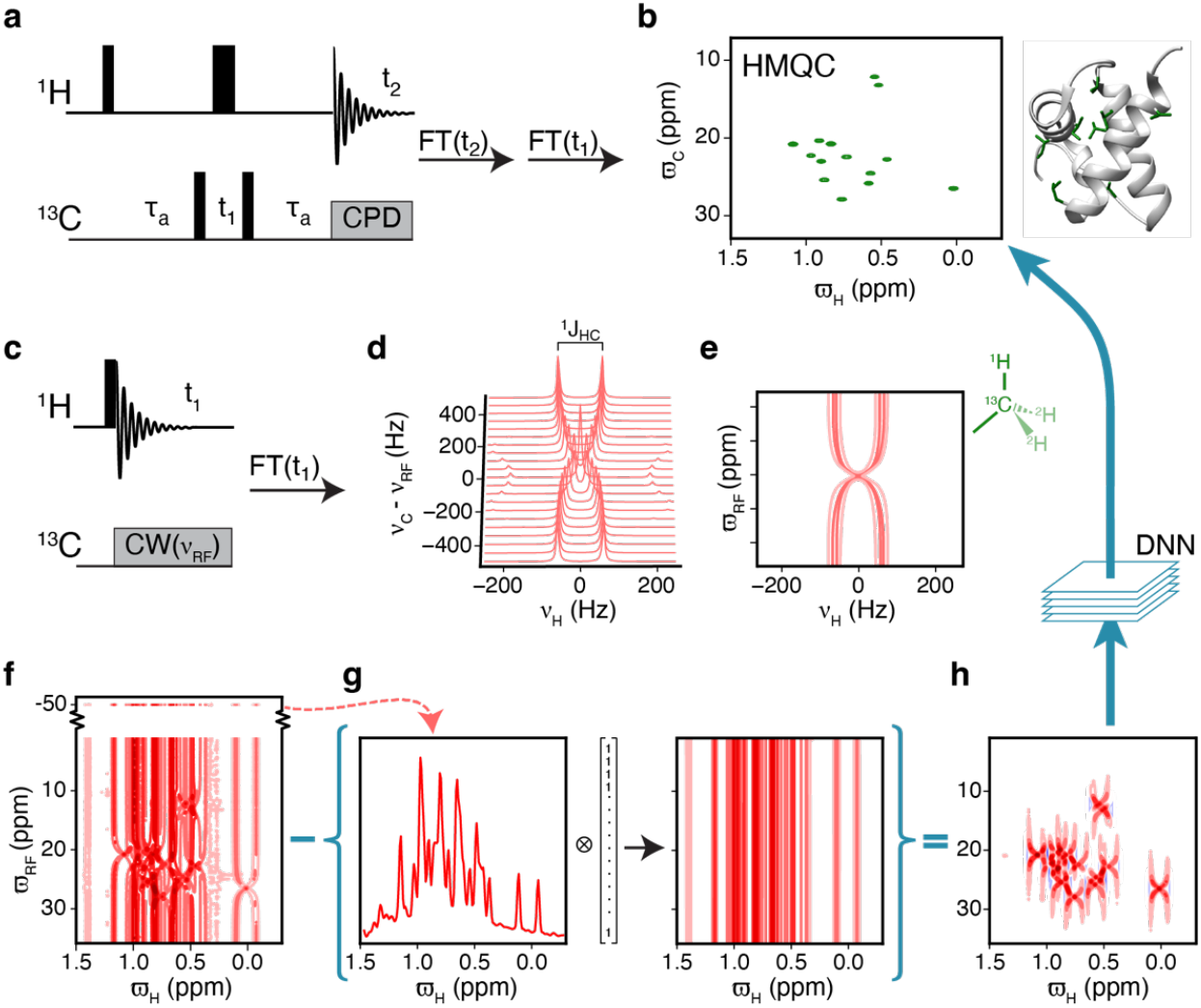
(**a**) The standard ^1^H-^13^C HMQC pulse sequence, which is traditionally used to obtain two-dimensional ^1^H-^13^C correlation spectra *via* 2D Fourier transformation. (**b**) The methyl region of the ^1^H-^13^C HMQC spectrum recorded on a ^13^CHD^2^ ILV isotopically labelled sample of the FF domain, which contains in total fourteen isoleucine, leucine and valine residues. (**c**) The ‘single pulse’ pulse experiment that forms the basis for the proposed approach to obtain 2D ^1^H-^13^C correlation maps. After the ^1^H excitation pulse, a weak ^13^C constant-wave (CW) decoupling is applied at various ^13^C offsets, *ν*_RF_, while recording the ^1^H spectrum. Fourier transform along the ^1^H dimension provides an off-resonance dataset as shown in (**d**) & (**e**) (contour plot). In (**d**,**e**) the off-resonance dataset is calculated using a ^13^C CW decoupling strength of *B*_1_ (^13^C) = 175 Hz and a one bond ^13^C-^1^H scalar coupling of ^1^*J*_HC_ = 125 Hz. The resonance frequencies of the ^1^H and ^13^C nuclei are set to 0 Hz. All the transverse relaxation rates were set to 25 s^-1^. (**f**) ^1^H-^13^C off-resonance dataset recorded on the FF domain, where substantial overlap is clear. (**g**) The 1D ^1^H reference spectrum of the FF domain, obtained with far off-resonance ^13^C decoupling, -50 ppm, contains doublets and solvent signals. (**h**) The ^1^H-^13^C (difference) off-resonance dataset obtained by subtracting the (coupled) ^1^H 1D reference spectrum in (**g**) from the ^1^H spectrum in (**f**). Overlap is reduced in the (difference) off-resonance dataset (**f** vs **h**) and can be transformed by a deep neural network (DNN) into a standard ^1^H-^13^C correlation map. All spectra of the FF domain were recorded on a 16.4 T (700 MHz) spectrometer at 6.5 °C.

## Results

Over the last couple of decades, it has become clear that methyl groups are sensitive reporters of the structure and dynamics of large proteins in solution. Consequently, several methyl NMR experiments and labelling strategies have been developed to study protein structure and dynamics (14). Concomitant with the discovery of the methyl-TROSY effect it was shown that methyl ^1^H-^13^C correlation maps with high S/N can be recorded using the simple four-pulse (^1^H-^13^C) HMQC experiment (33) (Fig. 1a,b) that exploits the methyl-TROSY effect (10). The HMQC experiment consists of two ^1^H↔^13^C transfer-periods (^1^H→^13^C and ^13^C→^1^H) and a ^13^C evolution period in addition to the ^1^H detection period (Fig 1a). About two decades later it was shown that an experiment, in which the ^13^C→^1^H transfer-period was eliminated by subsuming it into the ^1^H detection period provided spectra with substantially higher sensitivity for large proteins (15, 16). Datasets acquired using this experiment, referred to as delayed-decoupling HMQC (ddHMQC), are processed by the addition of a simple apodization prior to classical discrete Fourier transform. A natural step to attempt to obtain even higher sensitivity is to eliminate the last remaining transfer period (^1^H→^13^C) and consequently the ^13^C evolution period, which leads to the single-pulse experiment, Fig 1c. As detailed below, reconstructing ^1^H-^13^C correlation maps from such datasets is only possible due to the recent developments in machine learning and it is therefore now very timely to explore potential opportunities provided by the single-pulse experiment for characterizing large proteins.

### Brief overview of the single-pulse method

In the single-pulse experiment (Fig. 1c), following the single ^1^H (π/2) excitation pulse, the ^1^H free induction decay (FID) is recorded while applying ^13^C continuous wave (CW) decoupling (34, 35). The one dimensional (1D) ^1^H frequency-domain spectrum is obtained by Fourier transform of the FID and the process is repeated using different carrier offsets, *ν*_RF_, for the ^13^C CW decoupling. Fig. 1d illustrates the variation in the ^1^H spectrum as a function of the ^13^C carrier offset relative to the ^13^C carbon resonance frequency (*ν*_C_), *ν*_C_ − *ν*_RF_. In Fig. 1d *ν*_C_ = 0 Hz, the one bond ^1^H-^13^C coupling constant ^1^*J*_HC_, is 125 Hz and the ^13^C CW field is applied with a field strength of *B*_1_ = 175 Hz at various offsets, *ν*_RF_, ranging from −500 Hz to +500 Hz in steps of 50 Hz. A contour plot of the spectrum in Fig 1d is shown in Fig 1e, where an X shaped pattern is clearly visible. Overall, the ^1^H spectrum is approximately a doublet with an effective splitting of 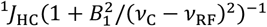(4, 35). It therefore depends on both the field strength of the ^13^C CW field, *B*_1_, as well as the carrier offset *ν*_RF_. Hence, the ^1^H spectrum consists of a doublet with peaks separated by ^1^*J*_HC_, when the ^13^C CW decoupling is carried out far from the resonance frequency of the ^13^C nucleus in question (|*ν*_C_ − *ν*_RF_| ≫ *B*_1_). On the other hand, when the carrier of the CW decoupling moves closer to the resonance frequency of the ^13^C spin, and |*ν*_C_ − *ν*_RF_| is decreased, the separation between the two peaks of the ^1^H doublet becomes smaller and the doublet collapses to a singlet when |*ν*_C_ − *ν*_RF_| approaches 0 Hz, giving rise to a unique X-shaped pattern centered at the ^13^C and ^1^H chemical shifts (Fig. 1e). Of particular interest here is that the series of ^1^H spectra recorded with ^13^C CW decoupling at different offsets, referred to here as an off-resonance dataset, contains information about the chemical shift of both the ^1^H (*ϖ*_H_; ppm) and the ^13^C (*ϖ*_C_; ppm) nuclei. In the past, off-resonance decoupling was used to correlate nuclei connected by ^1^*J* couplings, but these original studies were confined to small molecules due to spectral overlap (26-29, 36). As we aim to ultimately probe large proteins in solution, below we describe some of the strategies that we have used to alleviate the spectral overlap problem by leveraging recent advances in deep learning.

### Applying the single-pulse method to protein samples

Single-pulse off-resonance datasets of even small proteins lead to substantial overlap. The methyl ILV ^1^H-^13^C (HMQC) correlation map recorded using a sample of ^13^CHD_2_ enriched FF domain (∼8 kDa) contains a distinct peak for each of the fourteen methyl sites, Fig. 1b. The methyl ILV ^1^H-^13^C off-resonance dataset recorded on the same sample with *B*_1_ =175 Hz is shown in Fig. 1f. As expected from the calculations, the characteristic X-shaped patterns are observed, however, the off-resonance dataset is highly overlapped. One reason for the overlap in the off-resonance dataset are the streaks along the ^13^C dimension that arise due to the doublets separated by ^1^*J*_HC_. As mentioned earlier, these doublets arise when the ^13^C CW decoupling is carried out at an offset that is far from the resonance frequency of the ^13^C nucleus. To reduce overlap, a ^1^H 1D reference spectrum is recorded with far off-resonance ^13^C decoupling (-50 ppm; Fig 1f top), which is subtracted from every off-resonance 1D spectrum, Fig 1g. The reference 1D spectrum contains ^1^H doublets separated by ^1^*J*_*HC*_ and thus subtracting it from the off-resonance spectrum eliminates the streaks (Fig. 1h) and reduces the overlap (Figs. 1f *v.s*. 1h). Additionally, this subtraction also improves the baseline since strong solvent/buffer peaks etc. that are not affected by the ^13^C decoupling are also removed. A ^1^H spectrum recorded without ^13^C decoupling can also serve as the 1D reference spectrum. The off-resonance spectra from which the reference 1D spectrum has been subtracted will sometimes be referred to as difference off-resonance spectra.

Deep learning is renowned for extracting, simplifying and transforming features in complex datasets and has been extensively used to simplify and enhance NMR spectra (20, 37). Here we use a DNN to transform single-pulse off-resonance datasets (from protein samples) that contain the unique X-shaped patterns (features) into classical 2D NMR correlation maps with cross-peaks at the appropriate positions. This effectively means that a standard methyl 2D (^1^H-^13^C) correlation maps can be obtained using the single-pulse experiment that does not contain any ^1^H↔^13^C transfer periods or a ^13^C evolution period. In this proof of principle study we only focused on ^13^CHD_2_methyl groups because (*i*) the methyl region of a protein’s ^1^H-^13^C NMR spectrum contains peaks that originate only from methyl groups unlike for example the amide region that contain both ^15^NH and ^15^NH_2_ resonances, (*ii*) well established protocols are available to enrich proteins with isolated methyl groups within a silent deuterated background (14, 38), and (*iii*) to efficiently generate synthetic training data for the DNN one can treat the ^13^CHD_2_ groups as isolated two-spin (^1^H-^13^C) systems. To further aid the DNN in robustly transforming single-pulse off-resonance data, as discussed above, we provide the DNN with an additional pseudo-dimension consisting of two different off-resonance datasets, one with a *B*_1_ of 175 Hz and one with a *B*_1_ of 350 Hz. As we show below, this allows for the DNN to transform the two off-resonance spectra into the appropriate ^1^H-^13^C correlation map similar to the ^1^H-^13^C HMQC (or HSQC) (33, 39).

### Developing a DNN to transform off-resonance datasets into ^1^H-^13^C correlation maps

Training a DNN via supervised learning requires a large amount of training data, where the amount typically relates to the complexity of the neural network architecture (40). As this is a proof-of-principle study, we trained a simple DNN architecture with seven convolutional layers to carry out the transformation (see materials and methods, Fig. S2) as opposed to training the well-established but more complex FID-Net-2 (25, 41) that provides uncertainty estimates in addition to carrying out the desired transformation. The training data used to train the convolutional DNN consist of difference off-resonance datasets, akin to Fig 1h, and the corresponding target ^1^H-^13^C correlation map, akin to Fig 1b. As a large number of such datasets are not available, and DNNs have in the past been successfully trained to transform experimental magnetic resonance data using synthetic data (23, 42), we too generated synthetic data for training the DNN. More specifically, the difference off-resonance datasets consisted of two *I*(*ϖ*_H_, *ϖ*_C_) matrices (512×200) calculated for two *B*_1_ values (∼350 / ∼175 Hz). Synthetic off-resonance datasets were generated by propagating the Bloch equations with a range of parameters appropriate for methyl groups in proteins, including chemical shifts, relaxation rates, ^1^*J*_HC_ couplings, etc. Global parameters, those common to all methyl groups in one protein, include *B*_0_ (field strength ∼16.4 T), *B*_1_ (^13^C decoupling power, ∼350/175 Hz), ^13^C/^1^H sweep widths etc. were slightly varied while generating the training data (See materials and methods). The target ^1^H-^13^C correlation map was represented by a single *I*(*ϖ*_H_, *ϖ*_C_) matrix (512×200), with Gaussian-shaped peaks at the appropriate *ϖ*_H_ and *ϖ*_C_ chemical shifts. A Gaussian shape was used for the cross-peaks in the target, as opposed to a Lorentzian shape, because of its more localized nature and therefore a weaker requirement for the receptive field of the convolutional layers in the DNN. The linewidths, that is the Full-Width-at-Half-Maximum (FWHM), in the target ^1^H-^13^C correlation map of peaks in the ^1^H dimension is determined by *R*_2,H_/π, while linewidths of peaks in the ^13^C dimension were fixed to 30 Hz, as the off-resonance datasets do not directly provide ^13^C linewidth information. The DNN (Fig. S2) was trained to convert the synthetic input (∼350 / ∼175 Hz) difference off-resonance datasets into ^1^H-^13^C correlation maps (Fig. S3) by minimizing the mean square error (MSE) between the DNN reconstructed ^1^H-^13^C correlation map and the desired target ^1^H-^13^C correlation map. For an evaluation set of 250 synthetic datasets, containing a random number of peaks between 1 and 500, the MSE between the DNN reconstructed ^1^H-^13^C correlation map and the target ^1^H-^13^C correlation map was ∼1 compared to an MSE of ∼40 when an untrained DNN with random weights was used to reconstruct ^1^H-^13^C correlation maps. Two examples of how the trained DNN transforms (synthetic) difference off-resonance datasets into a ^1^H-^13^C correlation maps are shown in Fig. S3.

### Reconstructing experimental protein methyl ^1^H-^13^C correlation maps from off-resonance datasets

The trained DNN was first used to transform off-resonance datasets (Fig 2a) recorded on a sample of the ∼8 kDa FF domain, which, as mentioned earlier, contains fourteen ^13^CHD_2_ methyl groups. The DNN successfully transformed the off-resonance datasets (Fig. 2a) into a ^1^H-^13^C HMQC-like correlation map (Fig. 2b) in which peak positions agree well with those in the HMQC. In Fig. 2b, the DNN derived ^1^H-^13^C correlation map is shown in cyan while the standard HMQC ^1^H-^13^C map is shown in purple and an excellent agreement is observed. The peak positions along ^1^H and ^13^C, respectively, *ϖ*_H_ and *ϖ*_C_, obtained from the DNN derived ^1^H-^13^C correlation map agree well with those derived from the ^1^H-^13^C HMQC with RMSDs of ∼2 ppb and ∼26 ppb for ^1^H and ^13^C shifts, respectively (Fig 2e). We further proceeded to test the DNN using off-resonance datasets recorded on a sample of the medium sized T4L, which contains 60 (ILV) ^13^CHD_2_ groups (Fig. 2c). Although parts of the ^1^H-^13^C map are severely overlapped, the ^1^H-^13^C correlation map generated by the DNN (cyan) is very similar to the HMQC (purple) (Fig. 2d) and the chemical shifts of the peaks in the two spectra are again very similar, Fig. 2f, with RMSDs of 3 ppb and 23 ppb for ^1^H and ^13^C shifts, respectively.

**Fig. 2.**
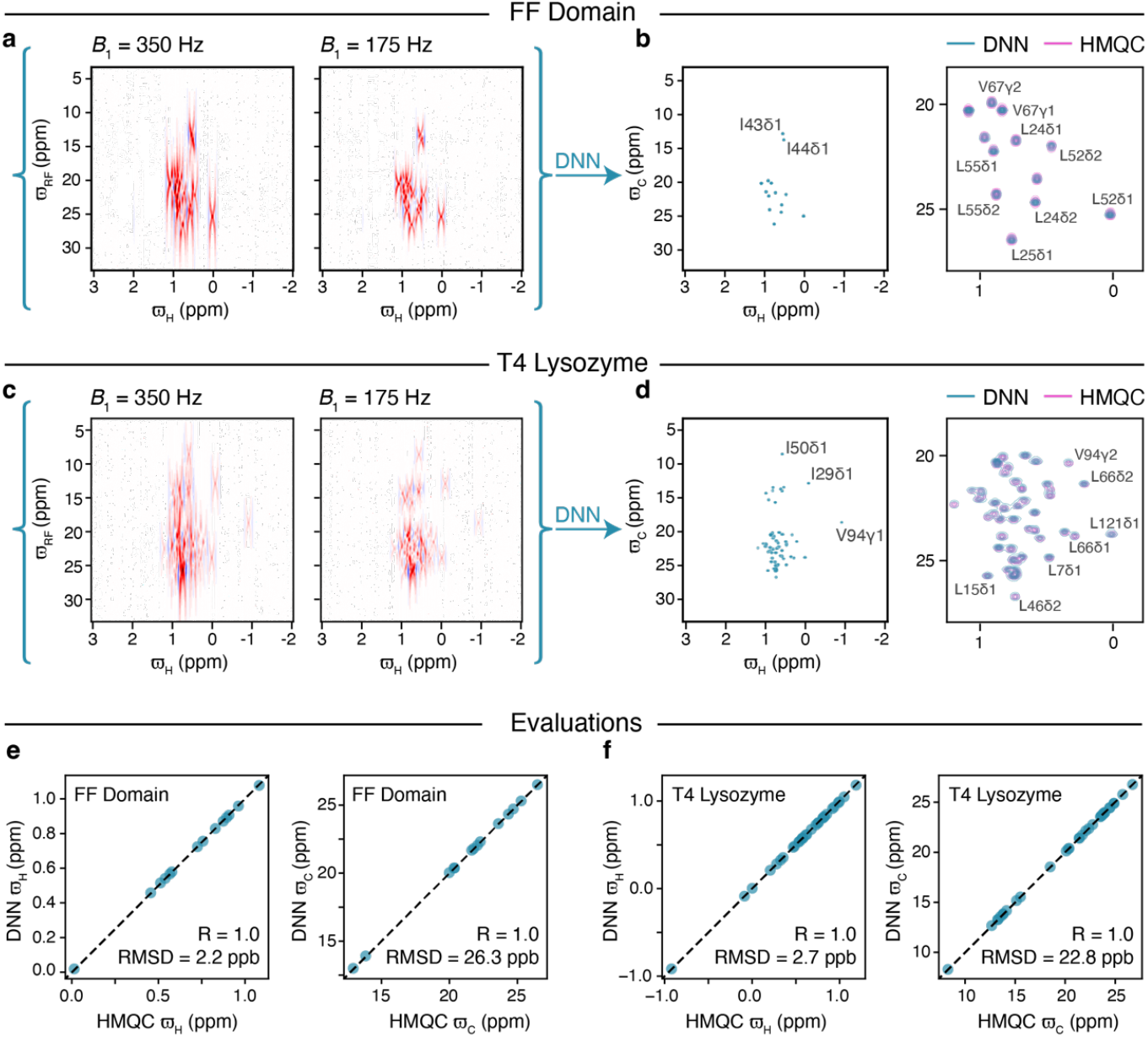
(**a**) Methyl ^1^H-^13^C difference off-resonance dataset for the FF domain with *B*_1_ (^13^C) = 350 Hz (left) and *B*_1_ (^13^C) = 175 Hz (right). (**b**) DNN reconstruction of the ^1^H-^13^C correlation map for the FF domain overlayed with the HMQC ^1^H-^13^C correlation map and along with a zoom of the most overlapped region (right). (**c**) Methyl ^1^H-^13^C difference off-resonance dataset for T4 Lysozyme with *B*_1_ (^13^C) = 350 Hz (left) and *B*_1_ (^13^C) = 175 Hz (right). (**d**) DNN reconstruction of the ^1^H-^13^C correlation map for T4 Lysozyme overlayed with the HMQC correlation map and along with a zoom of the most overlapped region (right). (**e, f**) Evaluation of accuracy of peak positions obtained from the DNN-derived ^1^H-^13^C correlation maps by comparison with peak positions from HMQC correlation spectra, shown for the FF domain (**e**) and T4 Lysozyme (**f**). All the experiments were carried out at 6.5 °C on a 16.4 T (700 MHz) spectrometer.

### Assessing the quantitative nature of the transformation

Having empirically established that the DNN transforms experimental data acquired using the single-pulse experiment into ^1^H-^13^C correlation maps with cross-peaks at the correct chemical shifts, we next set out to assess the quantitative aspects of the intensities of the cross-peaks in the transformed spectrum. Direct comparison between the intensities of peaks in the DNN generated ^1^H-^13^C correlation map and, for example, a ^1^H-^13^C HMQC/HSQC type spectra will not be meaningful as peak intensities in HSQC and HMQC spectra are sensitive to the ^1^*J*_HC_ couplings, ^13^C/^1^H relaxation rates, and to the exact values of the delays/evolution times used in the experiment. However, it will be meaningful to perform a comparison of relative peak intensities between DNN reconstructed ^1^H-^13^C correlation maps recorded using samples that are mixtures and in which the ratio of the constituents is varied. Thus, we constructed mixtures of the two proteins T4L and FF domain by combining off-resonance datasets from T4L and FF in ratios of 1:0.5 and 1:1.5, respectively (T4L concentration constant), and reconstructing the ^1^H-^13^C correlation maps using the trained DNN. The DNN derived ^1^H-^13^C correlation maps for the two ‘samples’ are shown in Fig. 3a and 3c and their uncertainties are shown in Fig 3b and 3d, respectively. The uncertainties were estimated by the Monte Carlo dropout method (see materials and methods).

**Fig. 3.**
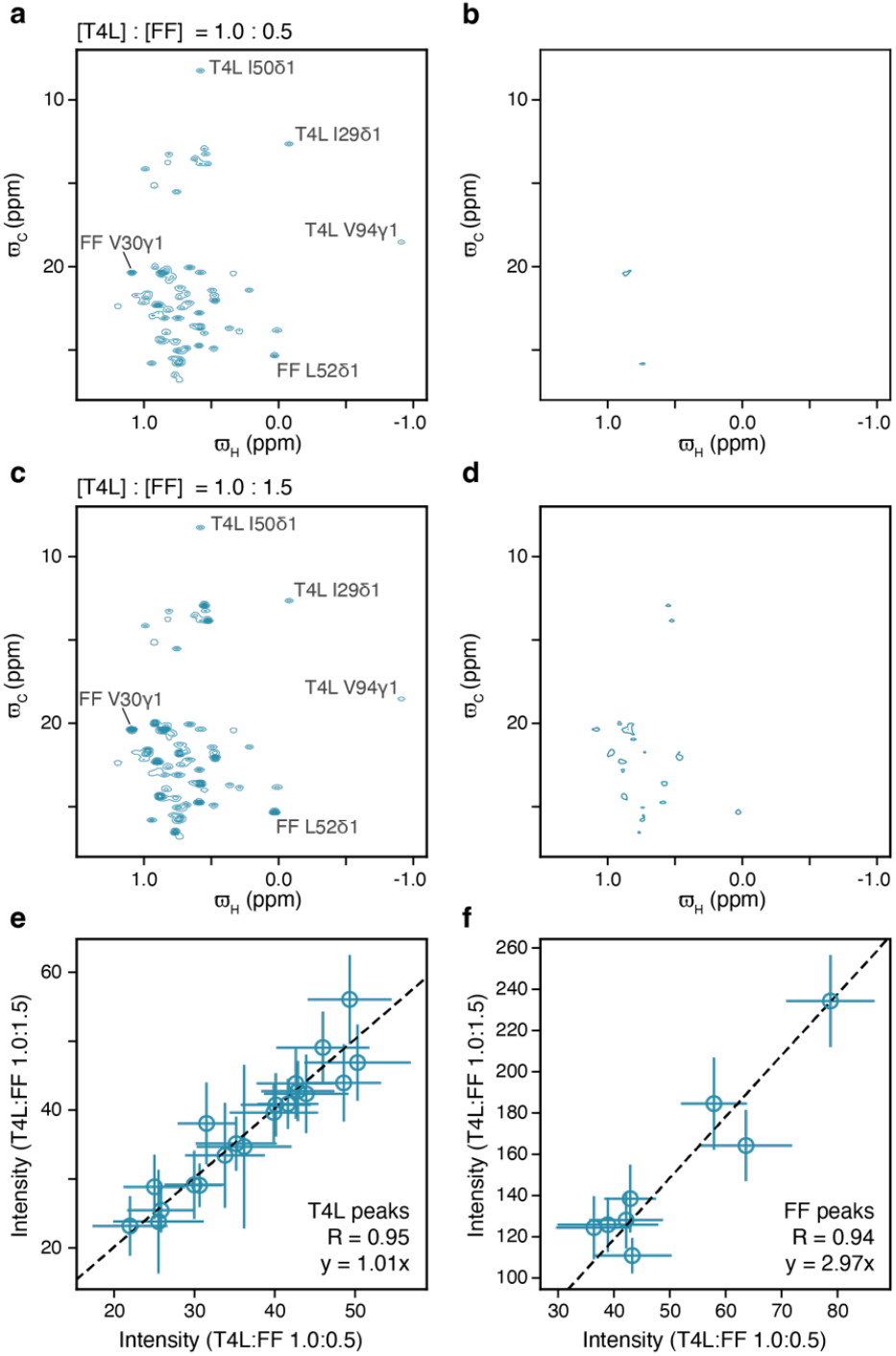
The peak intensities in the DNN reconstructed ^1^H-^13^C map report on concentrations of mixtures of the two proteins T4L and FF domain. DNN reconstructed ^1^H-^13^C maps (**a, c**) and associated intensity errors (**b, d**) when the T4L and FF domain difference off-resonance datasets are coadded in the ratio 1:0.5 (**a, b**) and 1:1.5 (**c, d**). Comparison of peak intensities between the DNN derived ^1^H-^13^C maps shown in **a** and **c** for resonances arising from T4L (**e**) and FF domain (**f**). Peak intensities are from data heights at the peak centers in **a** and **c** and the associated error bars are from **b** and **d**, respectively. Uncertainties (**b, d**) of the predicted intensities (*σ*_*Recon*_) in the ^1^H-^13^C correlation map were estimated using the Monte Carlo dropout procedure. Intensities arising from T4L peaks are identical within uncertainty in both the spectra as seen from the correlation in **e**, while the intensity of the peaks arising from FF domain increases by a factor of 2.97 close to the expected value of 3 (=1.5/0.5). The data used in the analysis presented here is from the experiments depicted in Fig. 2.

In line with the concentration of T4L being the same in both the samples, peaks that originate from T4L have similar intensities in Fig. 3a and 3c. On the other hand, peaks that originate from the FF domain are more intense in Fig. 3c (T4L:FF ratio of 1.0:1.5) than in Fig. 3a (T4L:FF ratio of 1.0:0.5) consistent with the concentration of the FF domain being higher in the sample corresponding to Fig. 3c. Fig. 3e compares the intensities of peaks that originate from T4L in the two DNN-derived ^1^H-^13^C correlation maps of the mixtures and, as expected, the intensities of the these peaks are similar in the two DNN derived spectra (slope of 1.01). Similarly, Fig. 3f compares the intensities of peaks that derive from FF in the two DNN derived ^1^H-^13^C correlation maps of the mixtures, and again as expected, the intensities of the FF peaks are higher the sample with ratio of 1:1.5, with a slope of 2.97 compared to the expected slope of 3 (1.5/0.5). Overall, this shows that the intensities of peaks in the ^1^H-^13^C correlation maps reconstructed by the DNN from experimental data are reasonably meaningful. It is possible that modifying the architecture of the DNN and a more informed choice of *B*_1_ fields used to record the off-resonance datasets will lead to better estimates of intensities and peak positions.

As the training data for the DNN did not include any artifacts, we wanted to assess the performance of the DNN in the presence of artifacts and thus access the DNN’s performance on slightly out-of-scope (out-of-distribution) data. Mis-setting the Z1 and Z2 shims on the NMR spectrometer, and thus altering the lineshape of the peaks in the input spectra, had almost no effect on the peak positions and also preserved relative peak intensities in the DNN generated FF domain methyl ^1^H-^13^C correlations maps (Fig. S4). While this does not include a full survey of all possible out-of-scope scenarios, it does show that, at least when it comes to lineshape, the DNN can reconstruct spectra under suboptimal conditions that were not included in the training data.

### Reconstructing the Ile ^1^H-^13^C correlation map of a 360 kDa protein

Having established that the single-pulse experiment, along with the trained DNN, can reconstruct the methyl ^1^H-^13^C correlation maps of small-to-medium sized proteins, we wanted to test the limits of the single-pulse strategy on larger systems with successively lower tumbling times. Thus, we prepared a sample of the ∼360 kDa double-α-ring (α_7_α_7_) of the proteasome that is ^13^CHD_2_ enriched at Ile δ1 positions. Single-pulse experiments were recorded at 50 °C and to make the task more challenging also at 10 °C. At 10 °C the viscosity of D_2_O is about 2.2 times higher than it is at 37 °C and about 2.95 times higher than at 50 °C (43). Thus, the relaxation properties of α_7_α_7_ at 10 °C are close to that of a ∼790 kDa sized protein at 37 °C and a 1 MDa particle at 50 °C.

The single-pulse/DNN derived methyl ^1^H-^13^C correlation map (50 °C) is in excellent agreement with both the ^1^H-^13^C HMQC spectra and the more sensitive delayed-decoupling ^1^H-^13^C HMQC, Fig. 4a, c, e. For the substantially more challenging case, α_7_α_7_ at 10 °C (Fig. 4b, d, f), several cross-peaks are missing and/or not well-resolved in the ^1^H-^13^C HMQC (Fig 4d) and ddHMQC (Fig. 4f) spectra. Thus, with this large molecule, the single-pulse/DNN derived correlation map (Fig 4b) appears to be qualitatively at par or slightly better than the HMQC & ddHMQC correlation maps (Fig 4d, f). For example, the peak indicated by an arrow (Fig 4b) in the 10 °C DNN derived correlation map is significantly broadened in the ^1^H-^13^C correlation maps (Figs 4d,f) derived using the HMQC/ddHMQC experiments. The HMQC and ddHMQC are very sensitive experiments for recording ^1^H-^13^C 2D correlation maps from ^13^CHD_2_ methyl groups in a _2_H background as the ^1^H-^13^C MQ coherence has very favorable relaxation properties because the ^13^C and ^1^H chemical shift anisotropies (CSAs) are small and the ^13^C-^1^H dipolar interactions essentially vanish. However, when evolution in the indirect dimension becomes unfavorable due to short transverse relaxation times, we expect that the single-pulse experiment will outperform traditional FT based 2D experiments for recording ^1^H-^13^C 2D correlation maps in large systems suggesting that it will be fruitful to explore the utility of the single-pulse strategy to record 2D correlation maps of very large macromolecular assemblies.

**Fig. 4.**
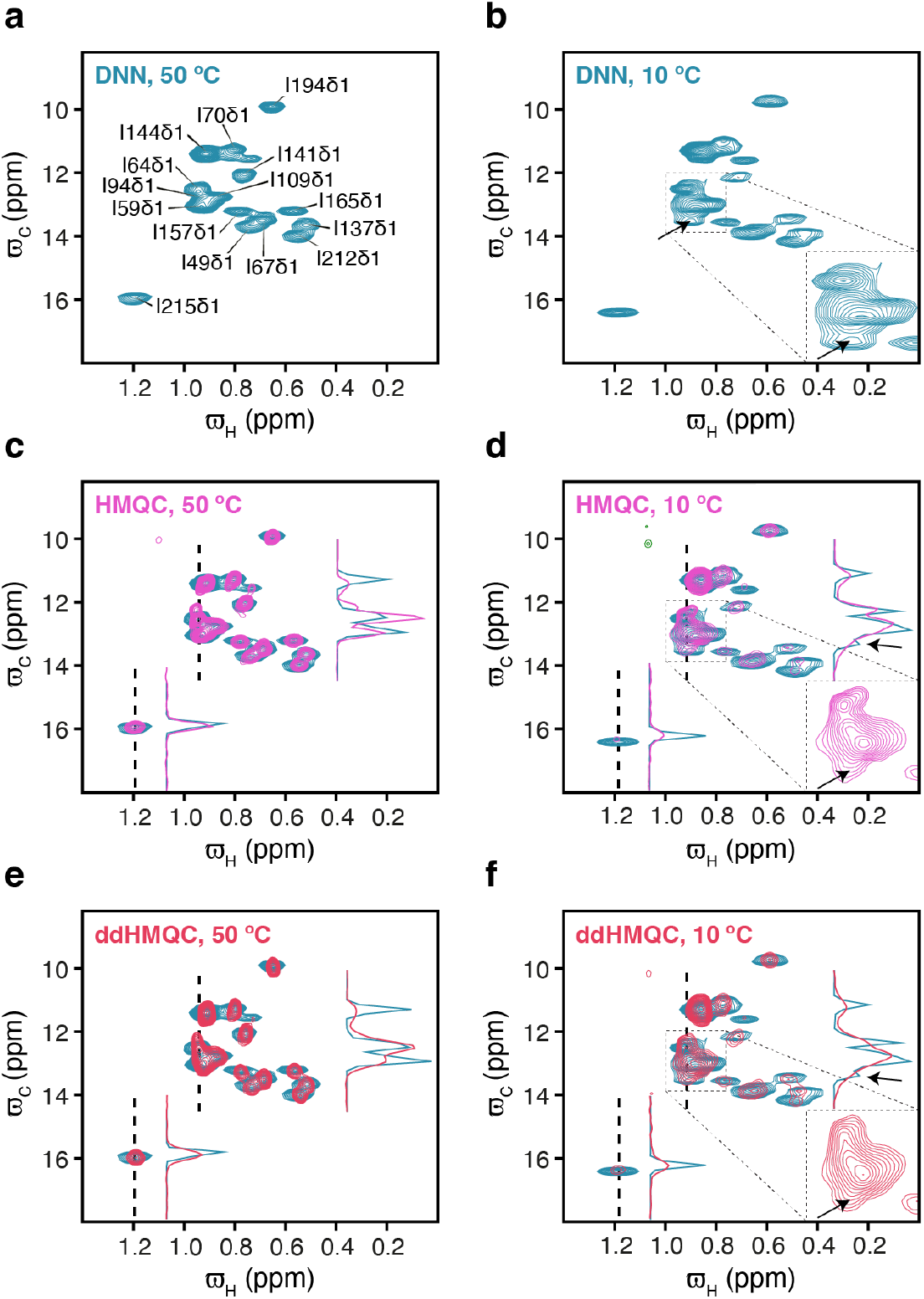
Ile *δ*1 ^13^C-^1^H correlation map (16.4 T; 700 MHz) of the *α*_0_*α*_0_ proteasome obtained at 50 °C (**a, c, e**) and 10 °C (**b, d, f**) using the DNN to reconstruct the correlation map from difference off-resonance data (**a, b**), a standard HMQC experiment (**c, d**), and a delayed-decoupling HMQC (ddHMQC) experiment (**e, f**). In (**c, d, e, f**) the DNN is shown in addition to the HMQC and ddHMQC. The HMQCs and ddHMQCs (**c, d, e, f**) were recorded with 32 scans, interscan delay of 1.5 s, acquisition times of 64 ms (^1^H) and 24 ms (^13^C, 48 complex points), and a total acquisition time of about 1.5 hours. The arrow in **b, d**, and **f** points to a peak that is resolved by the single-pulse experiment at 10 °C but is unresolved in the HMQC (**d**) and ddHMQC (**f)** spectra due to broadening in the ^13^C dimension. Representative 1D slices are shown to compare the HMQC and ddHMQC with the single-pulse experiment.

## Discussion

By developing a DNN to reconstruct the methyl ^1^H-^13^C correlation maps of both small and large proteins from off-resonance datasets obtained by single-pulse experiments, we have shown that the single-pulse experiment is not limited to small molecules but indeed, with the aid of deep learning it can provide high-resolution ^1^H-^13^C correlation maps of large protein complexes. In this proof-of-principle study we made use of ^13^CHD_2_ labelled proteins and obtained final ^1^H-^13^C correlation maps that are similar to standard HMQC (& ddHMQC) 2D correlation maps. However, the single-pulse approach has some shortcomings compared to the conventional FT approaches, such as the HMQC. In a conventional experiment all points in the indirect dimension contribute to the final signal in the frequency-domain spectrum (44) so long as the evolution time is not long compared to the transverse relaxation time *T*_2_. In contrast, in the single-pulse experiment presented here, only data recorded with ^13^C offsets, *ϖ*_RF_ close to the chemical shift of the ^13^C nucleus, *ϖ*_C_, contribute to the signal. Moreover, similar to other non-linear processing strategies, the transformation performed by the developed DNN, from single-pulse off-resonance datasets to 2D correlation maps, is not linear and therefore it does not transform gaussian noise into gaussian noise. Consequently, small artefacts might appear in the reconstructed spectrum and care must therefore be taken.

As this is a proof of principle study to show that the single-pulse/DNN strategy can generate 2D correlation maps, there is room for improvement and innovation. A relatively shallow and simple DNN was used because it allowed for fast training and for exploration of initial conditions. The limited capacity of the relatively small DNN used here can lead to higher (epistemic) uncertainties in the reconstructions. While we have estimated the uncertainties in the reconstructed maps using the Monte Carlo dropout procedure (see materials and methods and Fig. S3) (45) it is imperative for future applications to predict quantitative uncertainties as it will allow one to identify artefacts in the reconstructions. Larger architectures, e.g. FID-Net-2 (25), will result in both robust reconstructions over a broader range of input data and quantitative estimates of uncertainties. Furthermore, the optimum set of *B*_1_ fields and ^13^C offsets at which ^13^C CW decoupling is carried out has also not been determined. Optimizing these parameters, potentially aligned to the samples investigated and labeling schemes used, will likely improve the sensitivity and resolution of the resulting reconstructed 2D ^1^H-^13^C correlation map. It may also be fruitful to train DNNs that use regular HMQC/HSQC and off-resonance datasets to generate 2D correlation maps with higher resolution.

The single-pulse/DNN strategy was in this study demonstrated only for ^13^CHD_2_ methyl groups but the demonstration paves the way for applications encompassing other sites, such as more complicated spin-systems including ^13^CH_3_ methyl groups and aromatic groups in uniformly ^13^C enriched proteins and nucleic acids. One can also envisage the single-pulse/DNN strategy being extended to three-dimensional (3D) NMR experiments. In conclusion, this study not only illustrates the tremendous potential of the single-pulse/DNN strategy to obtain 2D NMR spectra from experiments without INEPT transfer periods, but it more generally again illustrates the tremendous potential of developing new NMR experiments together with customized DNNs. Specifically, the presented single-pulse/DNN strategy further adds to the emerging picture that optimal information can be extracted by developing DNNs to reconstruct readily interpretable NMR spectra from complex data that contains a large amount of information.

## Materials and Methods

### NMR Samples

The FF domain, T4L and the *α*_7_*α*_7_ half-proteasome were expressed in *E. coli* BL21(DE3) cells grown in 100% D_2_O M9 media containing 1g/L ^15^NH^4^Cl and 3g/L [U-^2^H] glucose. Appropriate precursors were added an hour before induction as detailed elsewhere (38). The FF domain (46), T4L (47, 48) and *α*_7_*α*_7_ half-proteasome (32) samples were purified as described previously. The FF domain sample contained ∼1 mM [U-^15^N, ^2^H], Ile δ1-[^13^CHD_2_], Leu, Val-[^13^CHD_2_,^12^CD_3_] protein dissolved in ∼550 μl 50 mM sodium acetate, 100 mM NaCl, 2 mM EDTA, 2 mM NaN_3_, pH 5.7, 100% D_2_O buffer. The T4L sample contained ∼1 mM [U-^15^N, ^2^H], Ile δ1-[^13^CHD_2_], Leu, Val-[^13^CHD_2_,^12^CD_3_] protein dissolved in 50 mM sodium phosphate, 25 mM NaCl, 2 mM NaN_3_, 2 mM EDTA, pH 5.5, 100% D_2_O buffer. The *α*_7_*α*_7_ half-proteasome sample contained ∼0.5 mM [U-^15^N, ^2^H], Ile δ1-[^13^CHD_2_] of the *α*-monomer dissolved in ∼550 μl of 20 mM sodium phosphate, 50 mM NaCl, 1 mM DTT, 1mM EDTA, pH 6.8 100% D_2_O buffer.

### NMR Experiments

All the NMR experiments on the FF (6.5 °C), T4L (6.5 °C) and *α*_7_*α*_7_ (10, 50 °C) samples were performed on 16.4 T (700 MHz) Burker Avance III HD spectrometers equipped with TCI cryoprobes. See Fig. S1 for additional details and the exact pulse sequence used to carry out the single-pulse experiment on ^13^CHD_2_ methyl groups. Complete off-resonances datasets at two *B*_1_ values were recorded in ∼1 (FF, T4L: 4 scans, interscan delay of 2s) or ∼1.5 (*α*_7_*α*_7_: 8 scans, interscan delay of 1.5s) hours.

### NMR Data analysis

NMRPipe (49) was used to process the NMR data and nmrglue (50) was used to transform the data between different formats. Keras (51) and Tensorflow (52) were used to both train the DNN and to construct ^1^H-^13^C correlation maps from the off-resonance datasets using the trained DNN. SPARKY (53, 54) was used to visualize the data. Peak intensities were quantified using SPARKY and PINT (55).

### Generating synthetic training data

Training data consisted of off-resonance datasets (array size: 512×200×2) as the input and the desired ^1^H-^13^C correlation maps (array size: 512×200) as the target. Both the off-resonance datasets and ^1^H-^13^C correlation maps consisted of 512 points in the ^1^H dimension and 200 points in the ^13^C dimension. The off-resonance datasets were calculated for two different *B*_*1*_ values (∼350, ∼175 Hz) by propagating the Bloch equations for a two-spin system. The Liouvillian (56) constructed using eight basis set elements (*H*_*x*_, *H*_*y*_, 2*H*_*x*_*C*_*x*_, 2*H*_*y*_*C*_*x*_, 2*H*_*x*_*C*_*y*_, 2*H*_*y*_*C*_*y*_, 2*H*_*x*_*C*_*z*_, 2*H*_*y*_*C*_*z*_) is given by:

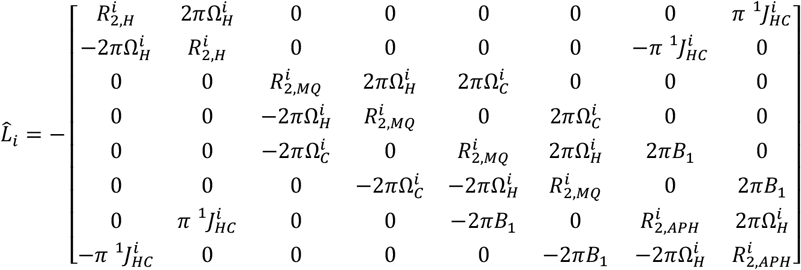

Here, 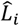 is the Liouvillian for ^1^H-^13^C spin-system *i*, leading to cross-peak *i*. Here, 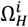 is the offset of the resonance frequency (Hz) of the ^1^H nucleus of peak *i* from the ^1^H carrier, 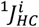 is the ^1^*J* coupling constant between the ^1^H and ^13^C nuclei of spin-system *i, B*_1_ (Hz) is the strength of the ^13^C decoupling field being applied along *x*, 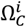 is the offset of the resonance frequency (Hz) of the ^13^C nucleus of *i* from the ^13^C carrier, 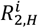 is transverse relaxation rate of the ^1^H nucleus, 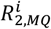 describes the relaxation of the multi-quantum terms (2*H*_*j*_*C*_*k*_, *j, k* ∈ (*x, y*)) of *i* and 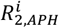 describes the relaxation of antiphase terms 2*H*_*x*_*C*_*z*_ and 2*H*_*y*_*C*_*z*_. Cross-correlated relaxation, which could lead to cross-relaxation between the different terms, was not considered. The FID originating from spin-system *i* is given by, 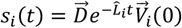 with 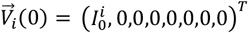 and 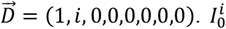 is the starting magnetization originating from *i*. The coupled (reference) FID, *s*_*u,i*_(*t*) is similarly calculated with *B*_1_ set to 0 in the above expression for 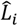.

To generate random off-resonance datasets for training and evaluation, experimental parameters (*B*_0_, *B*_1_, ^13^C sweepwidth, ^1^H sweepwidth, and phase of the FID) are randomly chosen over the ranges given in Table S1. The number of peaks (Npeaks) per spectrum is set to a random number between 1 and Npeaksmax (the maximum number of peaks in the spectrum). For each of the Npeaks peaks, peak specific Parameters 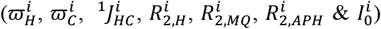 are randomly chosen over the ranges specified in Table S1. 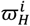 and 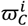 are respectively the chemical shifts (ppm) of the ^1^H and ^13^C nuclei of peak *i*. For a given ^13^C offset the peak (*i*) specific FID (*s*_*i*_(*t*)) is calculated for times varying from 0 to 64 ms in steps of 1/(^1^H sweepwidth) using 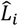 defined above. FIDs from all the peaks are summed to obtain the sample’s difference FID, 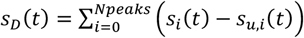. The sample FID is cosine square apodised, zero filled to 512 points and Fourier transformed to obtain the ^1^H spectrum while decoupling at a specific ^13^C offset using a specific *B*_1_ field. This process is repeated for all the desired offsets (200) and two *B*_1_ fields to generate the input off-resonance datasets. The off-resonance datasets were scaled so that the maximum value is unity and random gaussian noise with varying maximum values (see Table S1) was added following which the off-resonance datasets were rescaled so that the maximum value was unity. The corresponding training target, the ^1^H-^13^C correlation map, contains gaussian peaks at resonance positions of the peaks that were used to construct the off-resonance datasets. The linewidth of peaks in the ^1^H dimension is determined by *R*_2,*H*_ (with the minimum linewidth set to 20 Hz). On the other hand, the linewidth of peaks in the ^13^C dimension is fixed to 30 Hz. The intensity of the peaks is determined by the peak specific *I*_0_ values. Noise was not added to the training target.

### Training the DNN

The DNN used to transform the ^1^H-^13^C difference off-resonance datasets into the standard ^1^H-^13^C correlation is a convolutional neural network (57) with seven hidden layers (Fig. S2). The DNN was trained to transform the synthetic input off-resonance datasets (*B*_1_ ∼350, ∼175 Hz) into the desired output ^1^H-^13^C correlation maps using backpropagation via the ADAM optimization algorithm (58). During the training run, training datasets were continuously generated and never reused. The loss function was the mean square error between the intensities of the predicted and desired noise-free training target ^1^H-^13^C correlation maps. To reduce the chances of overfitting, dropout (30%) and L2 regularization were also used during training. At the start of training Npeaksmax was set to 25 and was gradually increased to a final value of 1500 (Fig. S3). The DNN was trained using ∼185,000 training spectra containing a total of ∼50 million peaks.

### Reconstructing the ^1^H-^13^C correlation maps from off-resonance datasets using the DNN

The experimental off-resonance (*B*_1_ ∼350, ∼175 Hz) datasets recorded at 200 ^13^C decoupling offsets and the reference ^1^H 1D spectrum were manipulated using NMRpipe to construct the difference off-resonance spectra (^1^H sweep width of ∼5 ppm) of the appropriate size (512, 200). The ^1^H FIDs were phased, cosine square apodised and appropriately zero filled as was done during training and so that the final spectrum has 512 points in the ^1^H dimension. Subsequently the data were Fourier transformed to generate the difference off-resonance spectra that were appropriately scaled, with a maximum value of 1, and used as the inputs to the DNN to construct the ^1^H-^13^C correlation map. Twenty ^1^H-^13^C correlation maps were predicted via a Monte Carlo dropout procedure (45) with 30% dropout in all the hidden layers. The mean of the twenty predicted *I*(*ϖ*_H_, *ϖ*_C_) maps was used as the reconstructed ^1^H-^13^C correlation map and as described in Fig. S3 the standard deviation, 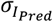, among the twenty predicted *I*(*ϖ*_H_, *ϖ*_C_ ) maps was used to estimate the point-by-point uncertainty, *σ*_*Recon*_, in the reconstructed correlation map with 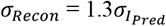.

## Software availability

Keras/TF scripts to carry out the reconstruction along with weights and example FF domain data are available online (https://github.com/hansenlab-ucl/DNN_single_pulse).

## Acknowledgements

We thank TIFR Hyderabad for funding (DAE, Government of India, Project No. RTI 4007), the TIFR Hyderabad NMR facility for a generous grant of spectrometer time, Dr. K Rao for maintaining the NMR facility, Prof. R Ramakrishnan (TIFRH) for computational facilities, Prof. V Agarwal (TIFRH) for providing isotopes, and Prof. Lewis Kay (Univ of Toronto) for useful discussions. The BBSRC (BB/R000255/1) is acknowledged for supporting the NMR facility at University College London. Access to ultra-high field NMR spectrometers was supported by the Francis Crick Institute through provision of access to the MRC Biomedical NMR Centre. The Francis Crick Institute receives its core funding from Cancer Research UK (FC001029), the UK Medical Research Council (FC001029), and the Wellcome Trust (FC001029). For the purpose of open access, the author has applied a Creative Commons Attribution (CC BY) licence to any Author Accepted Manuscript version arising. This research is supported by the UKRI and EPSRC (DFH; EP/X036782/1).

