## Supporting Information for "Two-dimensional NMR from a Single Pulse: Reconstructing Heteronuclear 2D spectra via off-resonance decoupling and Deep Neural Networks"

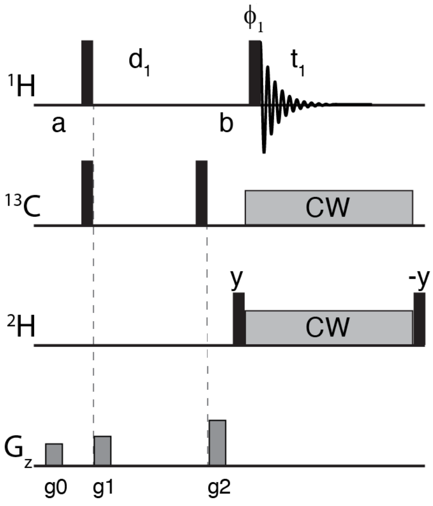


**Fig. S1.** 'Single Pulse' NMR experiment for recording 2D $^{1}H$-$^{13}C$ correlation maps from $^{13}CHD_2$ enriched protein samples. The pulses and gradients between points a and b are applied to destroy any unwanted coherences. Narrow black bars denote $\pi/2$ pulses and are of x phase unless indicated. $^{1}H$ and $^{13}C$ $\pi/2$ pulses are applied at the highest possible power while the $^{2}H$ $\pi/2$ pulses were ~150 μs. The $^{1}H$ carrier is placed on water, the $^{2}H$ carrier is placed at ~0.5 ppm in the middle of the methyl region. 0.5 kHz CW $^{2}H$ decoupling is carried out during $^{1}H$ detection. At point a, the $^{13}C$ carrier is placed at ~18.5 ppm in the middle of the methyl region and moved to the desired offset at point b. During detection, CW $^{13}C$ decoupling was carried out with $B_1$ values of ~350 or ~175 Hz. $\phi_1$, $\phi_{rec}$ = x, y, -x, -y. The $^{1}H$ FID was acquired for 64 ms ($t_1$). The gradients g0, g1 and g2 were applied for a duration of 300 μs with relative strengths of 0.35, 0.5 and 1.0 respectively. The experiment is carried out in a pseudo 2D manner with the $^{13}C$ CW decoupling carried out at different offsets. At 16.4 T (700 MHz) data was collected for a total of 200 $^{13}C$ offsets between -2636.75 and +2636.75 Hz around the reference carrier position (~18.5 ppm). A 1D $^{1}H$ spectrum without $^{13}C$ CW decoupling is also recorded. All the data was recorded with the $^{1}H$ sweep width set to ~16 ppm.

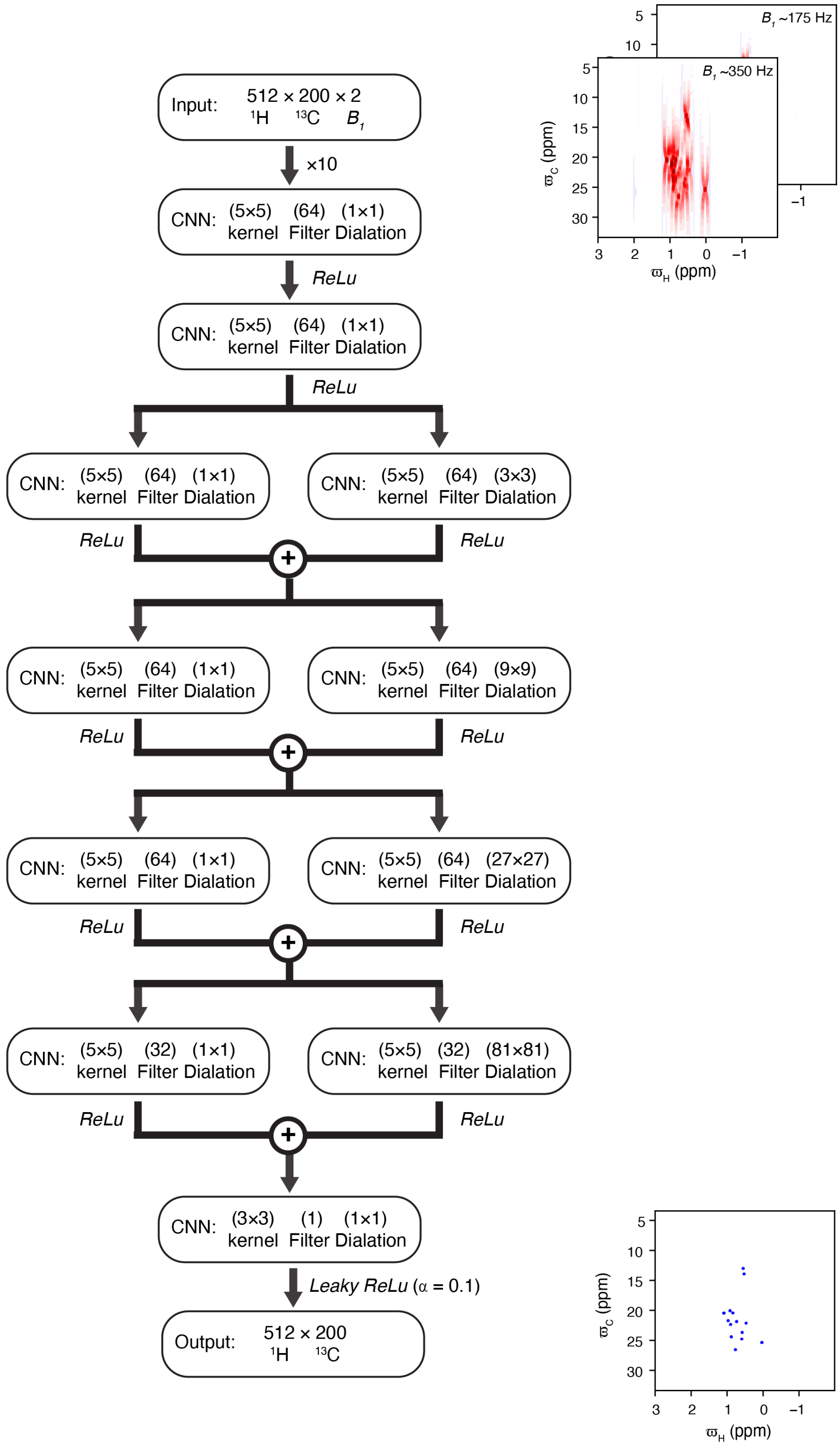


**Fig. S2.** The DNN used in this study to convert the ¹H-¹³C off-resonance data into a standard ¹H-¹³C correlation map consists of seven hidden layers. All the hidden layers are comprised of two dimensional convolutional neural networks (CNNs) whose parameters are indicated in boxes. Except for the last hidden layer, the activation function is the rectified linear unit (*ReLu*). The activation function for the last layer is a leaky rectified linear unit (*Leaky ReLu*). The + in the circle indicates concatenation. Padding is carried out so that the size of the output from every layer is 512×200 along the first two axis. The input consists of two ($B_1$ = ~350, ~175 Hz) difference off-resonance datasets. The input datasets are normalized by dividing the datasets by the largest intensity in both planes and internally multiplied by 10 (as indicated above) before being passed on to the first hidden layer.

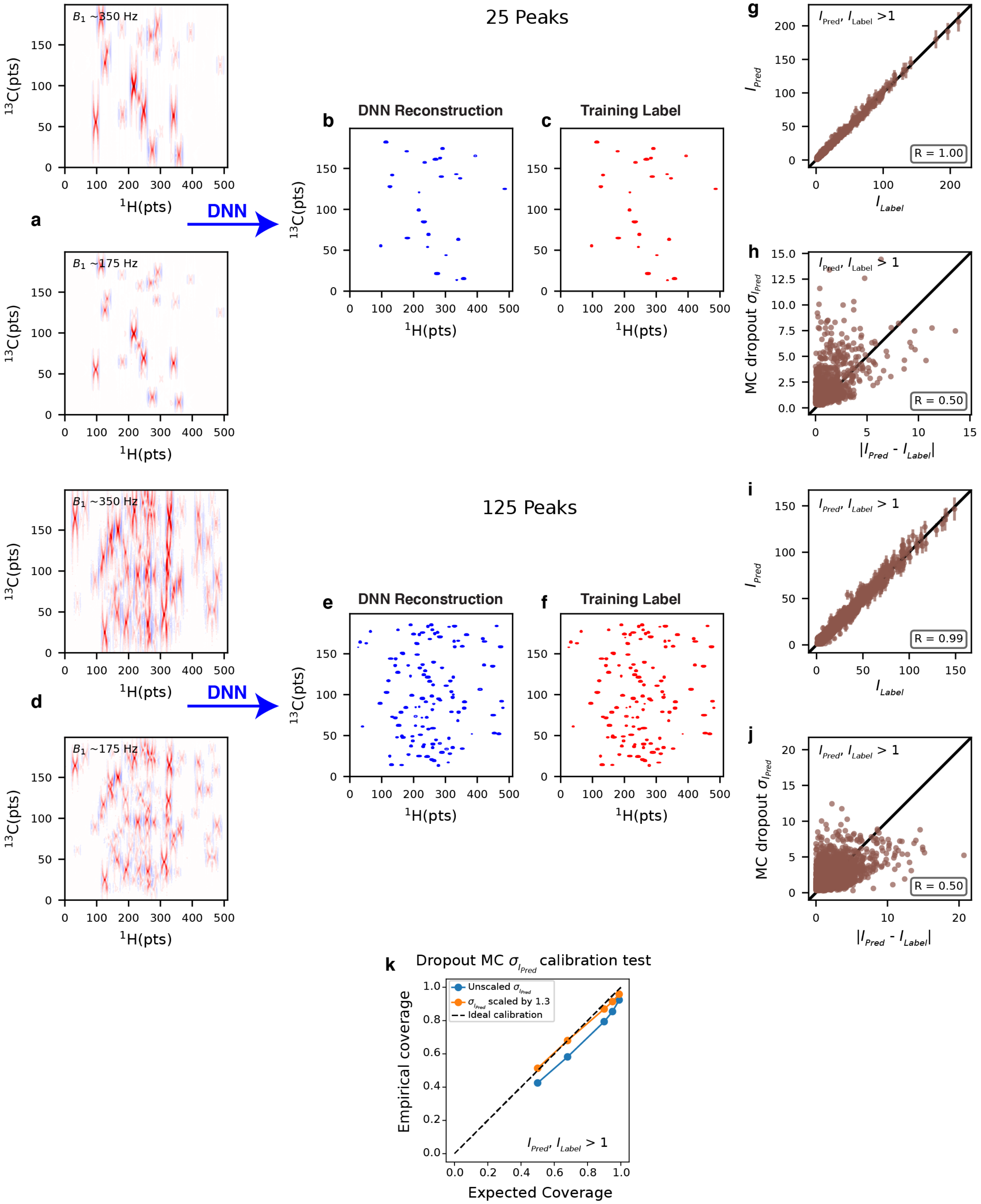


**Fig. S3.** Evaluating the DNN from Fig. S2 using simulated data. The DNN converts simulated difference off-resonance datasets with 25 (a) and 125 (d) correlations into 2D $^{1}$H-$^{13}$C correlation maps (b,e). For comparison the desired outputs (training label data) are shown in c (25 peaks) and d (125 peaks). A Monte Carlo dropout procedure with 20 trials was used obtain the reconstructed $^{1}$H-$^{13}$C correlation maps (See materials and methods). The mean of the 20 reconstructions is the DNN predicted ($I_{Pred}(\varpi_H, \varpi_C)$) $^{1}$H-$^{13}$C correlation map and $\sigma_{I_{Pred}}(\varpi_H, \varpi_C)$ is the point-by-point standard deviation of intensities among the twenty reconstructed maps.

(g,i) Comparison of the intensities ($I_{Pred}$) at various points of the DNN predicted spectra with intensities ($I_{Label}$) from the training label. In g (i) each point in b (e) is compared to the corresponding point in c (f). Comparison of (h,j) $\sigma_{I_{Pred}}$ with $|I_{Pred} - I_{Label}|$. The positive correlation suggests that the $\sigma_{I_{Pred}}$ values are meaningful estimates of the point-by-point intensity uncertainty ($\sigma_{Recon}$) in the reconstructed map. (k) Calibration test for the $\sigma_{I_{Pred}}$ values. If the $\sigma_{I_{Pred}}$ values are perfect estimates of the uncertainty, we expect ~68% of the $|I_{Pred} - I_{Label}|$ values to be ≤ $\sigma_{I_{Pred}}$ , ~95% of the $|I_{Pred} - I_{Label}|$ values to be ≤ $2\sigma_{I_{Pred}}$ and so on. The graph in k plots the fraction of $|I_{Pred} - I_{Label}|$ values empirically determined (empirical coverage) to lie within the expected value based on the $\sigma_{I_{Pred}}$ values versus the expected fraction (expected coverage). It is clear that the fraction of $|I_{Pred} - I_{Label}|$ values that lie within the expected range is reasonable but slightly underestimated based on the $\sigma_{I_{Pred}}$ values (compare blue and black curves). However, ~68% of the $|I_{Pred} - I_{Label}|$ values lie within $1.3\sigma_{I_{Pred}}$ and recalculating curve (orange) with scaled $\sigma_{I_{Pred}}$ leads to an excellent agreement (orange vs black) showing that the dropout MC derived $\sigma_{I_{Pred}}$ values are reasonable and that $1.3\sigma_{I_{Pred}}$ is a good estimate of the uncertainty ($\sigma_{Recon}$) in the reconstructed maps. Thus, in this study $1.3\sigma_{I_{Pred}}$ is used as an estimate of the uncertainty in the intensities of the DNN derived $^{1}H$-$^{13}C$ correlation maps. The plot shown in (k) was calculated using 250 difference off-resonance datasets, the corresponding $^{1}H$-$^{13}C$ correlation maps and DNN reconstructions. Each of the 250 datasets contained a random number of peaks between 1 and 500. In g-k only points in the correlation maps with $I_{Pred}$ and $I_{Label} > 1$ are considered.

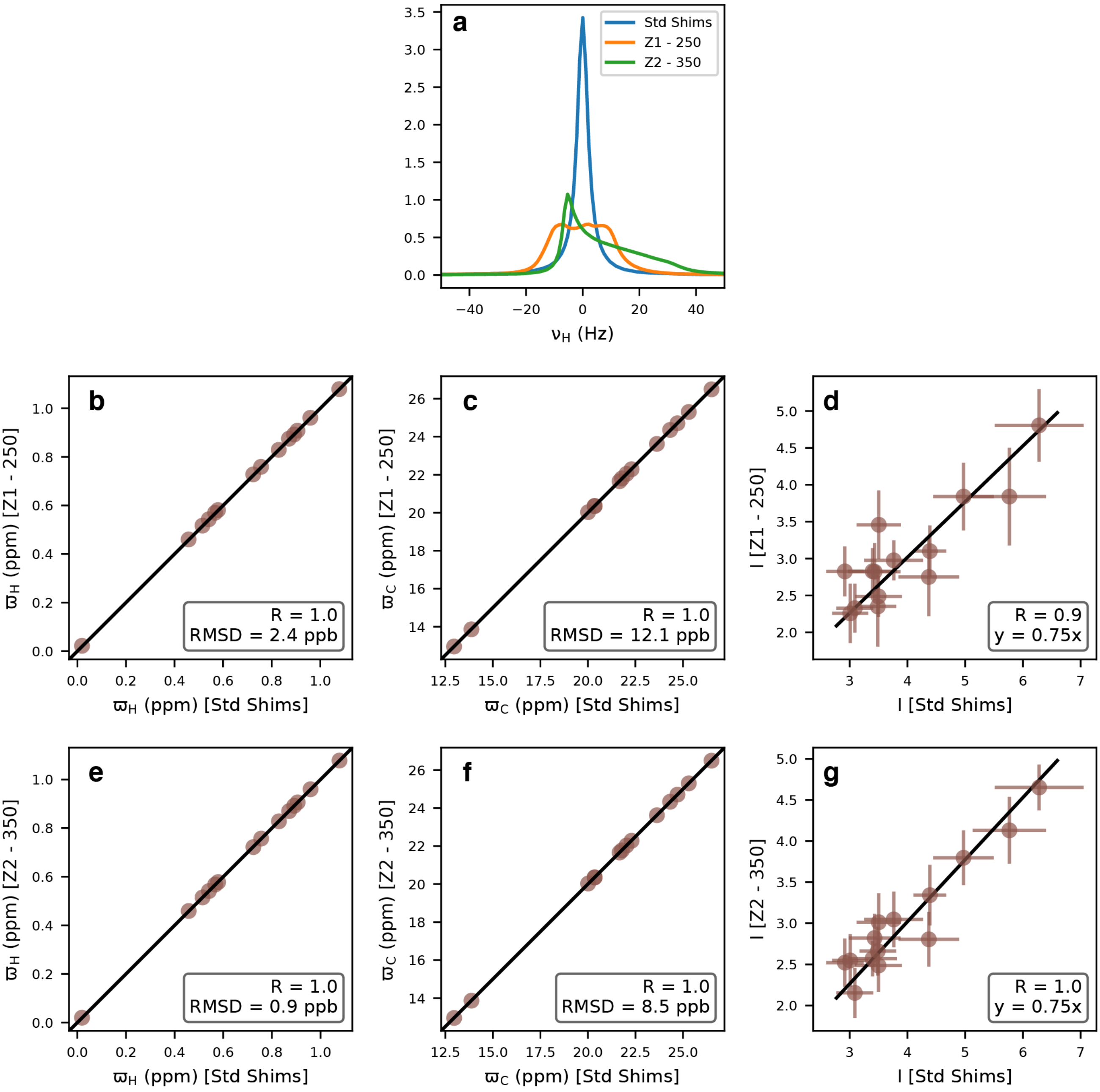


**Fig. S4.** Testing the robustness of the DNN by poorly shimming the magnet. a) Spectrum of the residual $^{1}$H signal in the FF sample with standard shims (Std Shims), Z1 decreased by 250 units (Z1 - 250) and Z2 decreased by 350 units (Z2 - 350). Comparison of $^{1}$H (b), $^{13}$C (c) peak positions and peak intensities (d) between the FF domain methyl ILV $^{1}$H-$^{13}$C maps constructed by the DNN with standard shims and with Z1 decreased by 250 units. Comparison of $^{1}$H (e), $^{13}$C (f) peak positions and peak intensities (g) between the FF domain methyl ILV $^{1}$H-$^{13}$C maps constructed by the DNN with standard shims and the Z2 decreased by 350 units.

| Spectral Parameters | | |
|---|---|---|
| **Parameter** | **Range** | **Comments** |
| $B_0$ | ± 0.5 MHz around the desired $B_0$ | For 700 MHz: 699.5 to 700.5 MHz |
| Number of Offsets (Noffset) | 200 [Fixed] | |
| Evolution time ($^1$H dimension) | 64 ms [Fixed] | |
| Number of points ($^1$H dimension) | 512 [Fixed] | |
| Spectrum Center ($^1$H,$^{13}$C) | (0 ppm, 17.5 ppm) [Fixed] | For convenience, does not matter |
| $^1$H sweep width | 4.85 to 5.15 ppm | |
| $^{13}$C sweep width | 29.0 to 31.0 ppm | |
| $B_1$ (First plane) | 325 to 375 Hz | $B_1$ does not have to be accurately calibrated. |
| $B_1$ (Second plane) | (0.48 to 0.52)× $B_1$ (First plane) | |
| Phase error | -5° to 5° | Phase error is introduced as the spectrum cannot be perfectly phased. |
| Gaussian error | 1% | Maximum noise in the input off-resonance datasets. |
| Number of Peaks (Npeaks) | 1 to 1500 | To include more overlapped examples during training. |
| **Peak Specific Parameters** | | |
| **Parameter** | **Range** | **Comments** |
| $^1$H chemical shift (ppm) | 1/3 peaks (-2.25 to 2.25 ppm)<br>2/3 peaks triangularly distributed around 0 between (-1.75 to 1.75 ppm) | Final range (-2.25 to 2.25 ppm) |
| $^{13}$C chemical shift (ppm) | ± 13 ppm | Final range (4.5 to 30.5 ppm) |
| $^1J_{H,C}$ | 115 to 140 Hz | |
| $R_{1,C}$ | 0.25 to 10 s$^{-1}$ | Internal parameter |
| $R_{2,C}$ | 1.25 to 125 s$^{-1}$ | Internal parameter |
| $R_{1,H}$ | 0.25 to 15 s$^{-1}$ | Internal parameter |
| $R_{2,H0}$ | 1.25 to 125 s$^{-1}$ | Internal parameter |
| $S1$ | 0.1 to 1.0 | Internal parameter |
| $S2$ | 0.1 to 0.75 | Internal parameter |
| $S3$ | 0.5 to 10.0 | Internal parameter |
| $R_{2,HEF}$ | $S1$ ×(1.25 to 125.0) | Internal parameter |
| $R_{2,H}$ | $R_{2,H0}$ + $R_{2,HEF}$ | Restricted to: 2.5 and 250 s$^{-1}$ |
| $R_{2,HE}$ | $R_{2,H}$ - $R_{2,H0}$ | Internal parameter, Restricted: 0 to 150 s$^{-1}$ |
| $R_{2,MQ}$ | $R_{2,HE} + S2 \times R_{2,C}$ | Restricted to: 1.25 and 250 s$^{-1}$ |
| $R_{2,APH}$ | $R_{2,H} + S3 \times R_{1,C}$ | |
| $I_0$ | 0.1 to 1.0 | To account for starting intensity variations due to varying $^1$H $R_1$s, sample mixtures with molecules of varying concentrations. |

**Table S1.** Parameters used to generate training data to train the DNN for reconstructing methyl $^1$H-$^{13}$C for off-resonance datasets. Unless specified '*x* to *y*' in the second column means that a uniformly distributed random real number *r* between *x* and *y* is chosen. In the third column 'Restricted to: *x* to *y*' means that if $r < x$, *r* is set to *x* and if $r > y$, *r* is set to *y*.
